# Oculomotor-related measures are predictive of reading acquisition in first grade early readers

**DOI:** 10.1101/2020.03.09.983197

**Authors:** Avi Portnoy, Sharon Gilaie-Dotan

## Abstract

Some estimates suggest that one in seven good readers and the majority of children with reading difficulties suffer from oculomotor dysfunction (OMD), an umbrella term for abnormalities in comfortable and accurate fixations, pursuits and saccades. However, national vision evaluation programs worldwide are often limited to distance visual acuity (dVA), not testing for OMD despite its high prevalence and the ease to detect it in brief optometric evaluations. We hypothesized that reading acquisition is dependent on good oculomotor functions and therefore inadequate oculomotor control will be associated with reading difficulties. We retrospectively examined and compared oculomotor evaluations and reading assessments (using standardized national reading norms) of a normative class (28 first graders (6-7 yr. olds)) that were independently obtained while blind to the other assessment. Almost a third (29.6%) were categorized as having OMD-related difficulties; better oculomotor performance was predictive of better reading performance. Control analysis revealed dVA was not positively associated with reading performance. Linear regression analyses further corroborated these findings. While this study is based on a small cohort and replications are needed, our results indicate that oculomotor functions are associated with reading acquisition and suggest that young children struggling with reading should be referred to a comprehensive visual evaluation including oculomotor assessments. This may allow providing treatments for learning-related visual issues.

## 2 Introduction

According to the Annie E Casey Foundation, in 2022, only 32% of children in fourth grade (9 year olds) in the US had reading skills that were proficient or above proficient level relative to their age (1). Sociologist Donald Hernandez found that the risk of leaving school without a diploma (awarded for the successful completion of high school in the USA) increases by 400% if a child enters their fourth year of formal education without proficient reading skills(2). Acquiring reading skills would therefore seem key to young adults finishing high school.

While reading is often considered a high-level perceptual and cognitive task involving language and comprehension, reading also involves and relies on the efficient functioning of both the visual motor and visual perceptual systems. Reading demands high-level binocular control, ensuring that the two eyes remain converged on the plane of the printed page whilst simultaneously shifting the eyes from word to word causing a relative asymmetrical adjustment in the angle of each eye. Similarly, the letters must be kept in focus, remaining sensitive to changes in accommodative demand as the eye shifts to lines closer or further from the reader. Accurate control of oculomotor functions such as fixation and saccades enable us to find and remain on the word currently being read, relative to the last word we have just read and the next word we are about to read. All these tremendously intricate and highly demanding oculomotor tasks are expected to take place with effortless automaticity simultaneously. These oculomotor functions take place concurrently with visual perceptual skills such as visual discrimination, visual memory and visual form perception etc. While for most good readers this is typically achieved with precision, this is not always the case for poor readers.

Oculomotor dysfunction (OMD) is an umbrella term for abnormalities in comfortable and accurate control of the oculomotor system with respect to maintaining fixation and/or eye movements (saccades and/or pursuits) (3,4). For those suffering from OMD, saccades can either overshoot (eyes land further than the designated target) or undershoot (eyes land short of the designated target). Both overshooting and undershooting are typically followed by smaller corrective saccades. In OMD saccadic onset latency may sometimes be increased and saccadic velocity may also be reduced. OMD may also be accompanied by unintentional motor responses to ocular movement (aka oculomotor overflow) of the jaw, tongue, head or body when planning or performing pursuits or saccadic movements. The extreme demands on the visual system that reading entails may incapacitate a new reader, with potential impact on both reading speed and accuracy. OMD’s prevalence in school-aged normative readers is estimated at 22-24% (5,6), compared to estimations as high as 96% in child populations diagnosed with a reading disability (5,7). It has been suggested that OMD may be linked to lack of effective integration of sensory input, higher-level decision-making, and appropriate oculomotor response (3,8,9).

There are studies supporting a connection between ocular control and reading ability (10–13), however in many European countries as well as in Israel and the USA, visual evaluations at early ages do not standardly assess oculomotor control and include only distance visual acuity (dVA) (14). We hypothesized that oculomotor skills, and particularly fixations and saccades, are critical for acquiring comfortable and efficient reading skills. To test our hypothesis and examine the relationship between reading acquisition and these oculomotor functions we retrospectively compared oculomotor results to independently obtained reading evaluations (Hebrew) in a normative class of first graders (6-7 year olds, n=28). For control, we also compared visual acuity assessment to the reading evaluations. The optometric evaluation was performed by an optometrist blind to the reading evaluation results; reading was evaluated by an experienced reading teacher blind to the optometrist’s evaluations.

## 4 Methods and Materials

### 4.1 Participants

The study includes retrospective analyses of data obtained from twenty-eight first grade children (6-7 year olds, 11 girls and 17 boys) that were all studying in the same class in a school that is part of the public education system in Israel.

The data from the optometric evaluations were obtained as part of the community outreach optometric evaluations activities routinely performed free of charge by the Bar-Ilan University School of Optometry and Vision Science (e.g. in schools, orphanages, centers for the elderly, immigration centers and additional community centers in rural and peripheral parts of Israel as well as in Bar-Ilan University Optometry clinics). The study protocol was in accordance with the guidelines of the institutional Bar-Ilan University ethics committee that approved retrospective analyses of data (stripped of identifying information) collected in optometric evaluations as well as scholastics evaluations to investigate the relationship between visual functions and scholastic performance.

The parents of the children, all in the same class, were offered the opportunity for their children to undergo an optometric evaluation in school and were asked to provide their consent in writing. Out of a class of thirty children, all but parents of two children provided written informed consent that their child participate in the optometric evaluation. The visual function evaluation was performed by a certified optometrist and took roughly twenty to thirty minutes per child. Children who already had a prior refractive correction wore their current daily glasses for each of the optometric tests. Parents were notified about any suspicious finding regarding their child’s vision that may require further evaluations. If visual acuity was found to be 6/12 or worse in either eye with their current spectacle correction (n=4), after the visual assessment the child was referred to an optometrist for further refractive testing or an ophthalmologist for a full ocular health exam (based on retinoscopy results).

In addition, as part of the school program, all the children underwent a comprehensive reading examination including reading speed and accuracy, performed by a reading teacher a couple of weeks after the optometric evaluation. Importantly, the teacher was not informed of the optometric results until a few weeks after the whole class’s reading assessments were completed, and the optometrist was not informed of the reading assessment results until after the optometric evaluations were completed. One child was excluded from the analysis due to significant language and learning difficulties observed independently in the optometric and reading evaluations. Another child’s dVA records were missing and we therefore had to remove him from the dVA related analyses.

### 4.2 Visual evaluation

The visual function evaluation was performed as part of the community outreach optometric evaluation activities the Bar-Ilan University School of Optometry and Vision Science engages in (see details above). Each child’s optometric evaluation included the following standardized optometric tests carried out in this order (more details below): Distance and Near visual acuity (VA) (15), objective refraction – retinoscopy (15), cover testing (distance and near) (15), negative fusional vergence (NFV) at near (prism bar) (15), positive fusional vergence (PFV) at near (prism bar) (15), near point of convergence (NPC) (15), stereopsis (16), accommodative amplitude (17) (right eye (RE) only (18)), accommodative facility (15) (RE only (19)), NSUCO (4), King Devick (20,21) and Developmental Eye Movement test (DEM) (22,23). Lighting was provided by overhead fluorescent fixtures as well as sunlight from windows. For the dVA testing (because a projector was used), the fluorescent lighting was extinguished, and the sunlight was partially blocked by curtains leaving the room moderately lit (24).

***Distance visual acuity*** (VA) was performed using a standard Snellen acuity chart using numerical optotypes projected onto a wall 6 meters from the child and performed both monocularly and binocularly (15). Since reading is performed with both eyes, the binocular distance VA measurement was used in the analyses of this study.

***Near VA*** was assessed at 40cm monocularly and binocularly using a standard near VA test card with numerical optotypes (15) (Rosenbaum pocket vision screener).

***Retinoscopy*** is an objective assessment of the child’s refractive status. This was measured to ensure that children not wearing glasses were not in need of any refractive correction and that children wearing glasses were not under or over corrected by more than 0.5 diopters (D) of myopia (short sightedness), 1.5D hyperopia (far sightedness) or 0.75D astigmatism(19). If a discrepancy equal to or above these refractive errors was found, they were referred to get corrective spectacles (n=4).

***Developmental Eye Movment (DEM) test*** was one of the tests used to assess oculomotor function and has vertical and horizontal subtests. The DEM is composed of digits organized in vertical or horizontal alignments (each arrangement in a different subtest, vertical assumed to predominantly rely on vertical saccades, horizontal assumed to rely predominantly on horizontal saccades) that the examinee has to read aloud and the scores are given according to speed and accuracy performance in each of these subtests. The time recorded for each subtest is adjusted based on the number of digits that were skipped or added by the examinee. An overall categorization of performance is then defined using the ratio of horizontal test speed divided by vertical test speed (assumed to be related to language related processes) where the purpose of the ratio score is to differentiate language automaticity difficulties (rapid automatized naming (RAN)) (25) from oculomotor difficulties (26), as explained in the DEM manual (27). Results are categorized into one of the four following types (26,27):

1. Normal language automaticity and oculomotor function.
2. Normal language automaticity with oculomotor dysfunction.
3. Abnormal language automaticity function but normal oculomotor function.
4. A combination of both oculomotor and language automaticity dysfunctions.

DEM speed and accuracy raw scores are standardized according to age or grade (grade used in this study, see individual data at https://osf.io/shw2v/) with a mean score of 100 and standard deviation of 15.

***NSUCO (Northeastern State University College of Optometry)/Maples*** oculomotor test was carried out according to the manual supplied by the Optometric Extension Program (OEP) (9). This test is nonverbal, testing both saccades and pursuits individually. The examinee is asked to follow targets that are held at different specific locations or moved according to specific spatial trajectories by the optometrist. Targets are round, reflective and approximately 0.5cm in diameter. Targets are positioned between 40cm and Harmon distance (distance from elbow to middle knuckle) of the examinee. The distance between targets or diameter of pursuits should be 10cm in each direction from the examinee’s midline (nose). This test grades eye movement (saccade and pursuit) ability, accuracy, and head movement and body movement accompanying the eye movements.

Each of the sub-scores is graded from 1 (poor performance) to 5 (good performance). Two sub-scores titled ‘head movement’ and ‘body movement’ are graded by how much head or body movement is triggered by ocular movement. This indicates an inability to dissociate ocular movement from head or body movement, which is a typical sign of OMD (9). A grade of 1 represents gross head or body movement (poor performance) and a grade of 5 represents no head or body movement (good performance). More detailed explanations of the scoring can be found elsewhere (9).

### 4.3 Reading assessment

Reading of each child was tested by an experienced reading teacher. The teacher used the age-based norm-referenced tests in Hebrew provided by the Israeli Ministry of Education (28) (see more details at https://cms.education.gov.il/EducationCMS/Units/Rama/AarachaBeitSifrit/Mivdak_Hebrew.htm) and followed their fixed protocol for testing. The assessment (29) comprised 10 different subtests assessing reading and writing skills including accuracy, speed, phonological awareness, and reading comprehension.

In our retrospective analysis we chose to focus on subtest 8 which assesses reading speed and accuracy (but not comprehension) since it best simulates visual demands of daily classroom reading. The text is a 77-word story presented in 8 widely spaced lines. The child is required to read it aloud while the teacher evaluates both accuracy and speed. The teacher marks all the words incorrectly read by the child including self-corrections. According to the national standards (28), a score of 69-77 words read correctly is categorized as a ‘pass’, a score of 58-68 words read correctly as ‘demands follow up’ and a score of or below 57 words read correctly as a ‘fail’. With regards to reading speed the teacher measures the amount of time it takes the child to read the 77 words. A score of 195 seconds to read the text or less is considered a ‘pass’, a score of 196 to 265 seconds is considered ‘demands follow up’ and a score of 266 seconds or slower is deemed a ‘fail’ (28).

### 4.4 Analyses

For each child the measured DEM speed (duration in sec) and the number of errors for each of the tests (A, B, C – two vertical and one horizontal) were inserted into the DEM Scorer Optometric Clinical Software Version 2.2 (© 2008 Software in Motion) that converted each measure to standardized scores (according to the child’s grade level following the guidelines provided in the DEM manual (27)). The corresponding standard scores, DEM type categorization, and the DEM ratio were calculated by the program (data available at https://osf.io/shw2v). Spearman rank order correlation analyses between reading performance (speed, accuracy) and visual functions (DEM standardized speed or accuracy and dVA) were run to evaluate whether there was a monotonic relationship between reading and a specific visual function. Comparisons of reading scores of the poorer visual performers and those of the better visual performers were also performed: for each visual function, the class was divided into two groups based on their visual performance (a group of higher visual scores and a group of lower visual scores). The reading scores of the two groups were then subjected to a non-paired, one-tailed t-test (unequal variance; the t.test function in R was used and this function relies on the Welch (or Satterthwaite) approximation for to the degrees of freedom for t-tests with unequal variance (more details on the R statistical package we used appears below)). We also present the distribution of the reading test outcomes (pass, follow up, or fail) for the better and poorer visual function groups. Because our cohort size contained an odd number of children (n=27), after sorting the class according to a specific visual function, the 14^th^ child was removed prior to analyzing the data. This enabled us to have two groups of equal size. Lastly, we ran a set of linear regression analyses to assess whether reading performance was predicted by DEM speed, accuracy, type, and/or dVA. For one child, data for dVA was missing and we therefore removed this child’s data for all the linear regression models so we could also have a multifactorial model.

All of the above analyses were performed with R Studio (RStudio 2022.02.3 Build 492, © 2009-2022 RStudio, PBC) running with R version 4.2.1 (released on 2022-06-2, aka Funny-Looking Kid, Copyright © 2022 The R Foundation for Statistical Computing, Platform: x86_64-w64-mingw32/x64 (64-bit)). See https://osf.io/shw2v/ for more detailed data and additional supplementary information.

## 5 Results

Based on years of clinical observations we hypothesized that reading acquisition would be highly dependent on oculomotor functions and therefore examined the relationship between OMD-related measures and reading acquisition scores in a normative group of first graders.

### 5.1 Oculomotor evaluation by DEM and the relation to reading performance

The developmental eye movement test (DEM) is a well-established quick optometric evaluation tool whose last section (test C with horizontal arrangement of numbers) is considered to mimic the saccadic demands of reading on the visual system (22,23,26,30–32). We therefore expected that DEM performance on horizontal test C would be predictive of reading performance during reading acquisition period in either speed, accuracy, or both.

We started by examining the relationship between DEM horizontal speed and reading speed. To that end we first correlated (Spearman rank correlation) DEM speed (standardized scores) with reading speed and found DEM speed to be significantly predictive of reading speed (horizontal *r_s_*(*25*)*=-.43, p=*.012) as we hypothesized (analyses with DEM raw speed scores were almost identical). In addition, we also compared the reading speed scores of slower DEM performers to those of faster DEM performers and found (Fig. 1A) that the faster DEM group (as determined by the horizontal test C speed (standardized scores), in grey in Figure 1A) were significantly faster on the reading test than the slower DEM performers (in orange, *t*(18.54)=2.63, *p*=.008, 1- tailed). In addition, as can be seen in Figure 1B, in the faster DEM group (test C) the proportion of ‘pass’ grades in reading speed was high (>80%) with no ‘fail’ (Fig. 1B) while in the slower DEM group only about a third (38.5%) received ‘pass’ in reading speed and 30.8% ‘fail’. The results of the vertical DEM test were similar and are presented in Fig. 1C and D (correlation of DEM vertical speed and reading speed: *r_s_*(25)=-.51, *p*=.003; reading speed of faster vs slower DEM vertical groups: *t*(18.53)=2.76, *p*=.006, 1-tailed; faster DEM group: 92% ‘pass’ in reading speed and 0% ‘fail’, slower DEM group: 30.8% passed, 23.1% failed).

**Fig. 1.**
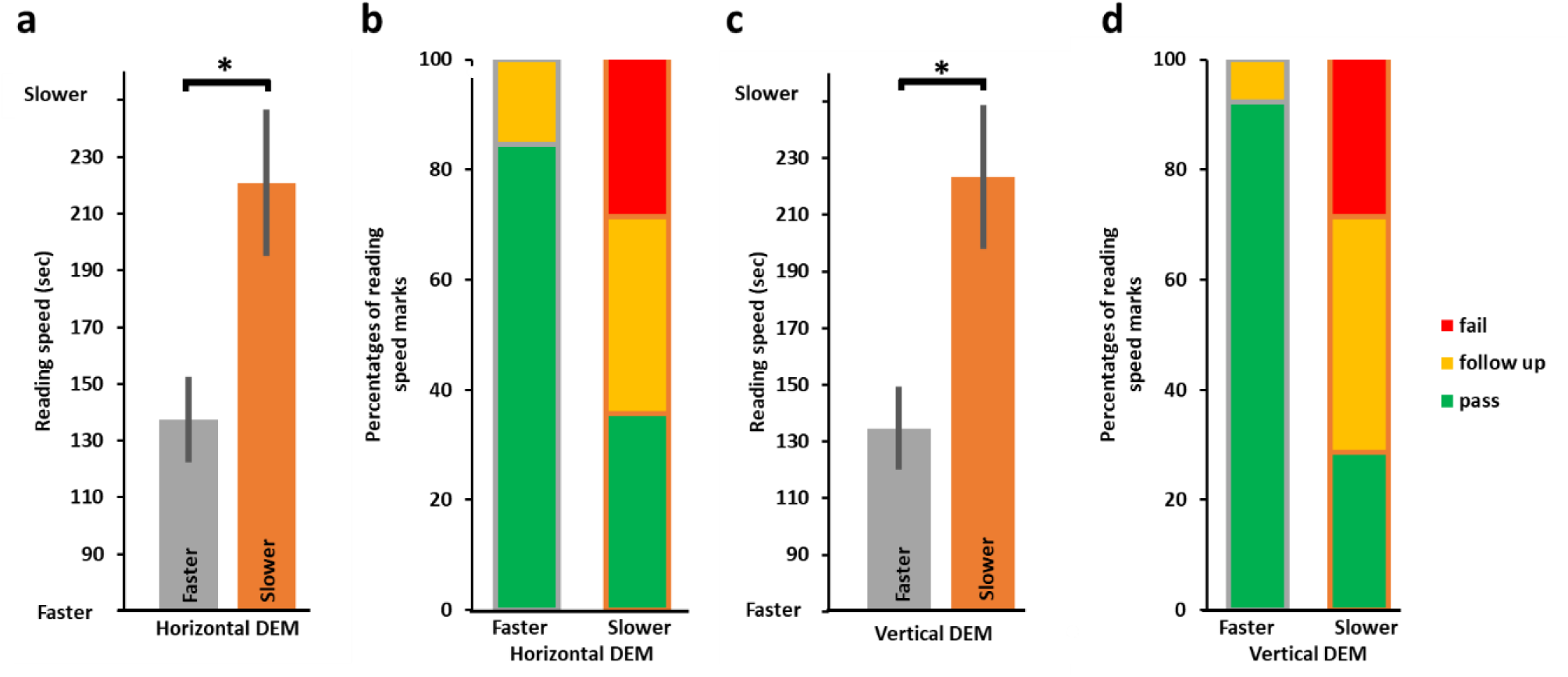
Faster reading speed and better reading grades for faster DEM performers (n=26). **(A)** Average reading time in seconds for slower (orange, on right, n=13) and faster (grey, on left, n=13) horizontal DEM performers; the faster horizontal DEM performers were found to be significantly faster in their reading performance (*t*(18.54)=2.63, *p*=.008, 1- tailed). **(B)** Proportions of reading speed categorization (pass in green, follow up in yellow or fail in red) for the slower DEM performers (n=13, right) and the faster DEM performers (n=13, left). **(C, D)** Same analysis as in (A, B) but for vertical DEM speed, the faster vertical DEM performers were found to be significantly faster in their reading performance (*t*(18.53)=2.76, *p*=.006, 1-tailed). Note that there were no fails for the reading speed test in both the faster vertical and faster horizontal DEM groups. Asterisks denotes *p*<.01.

Next, we examined the relation between DEM accuracy and reading accuracy. DEM horizontal accuracy and reading accuracy were not correlated (*r_s_*(25)=.18, *p*=.179). The more accurate DEM group were on average more accurate on their reading but this did not reach significance (*t*(16.26)=-1.31, *p*=.104, 1-tailed, see Fig. 2A). This was reflected in better reading grades (see Fig. 2B, the more accurate DEM group with 69.2% passes in reading accuracy and only 15.4% fails, the less accurate DEM group with 53.8% passes and 30.8% fails).

**Fig. 2.**
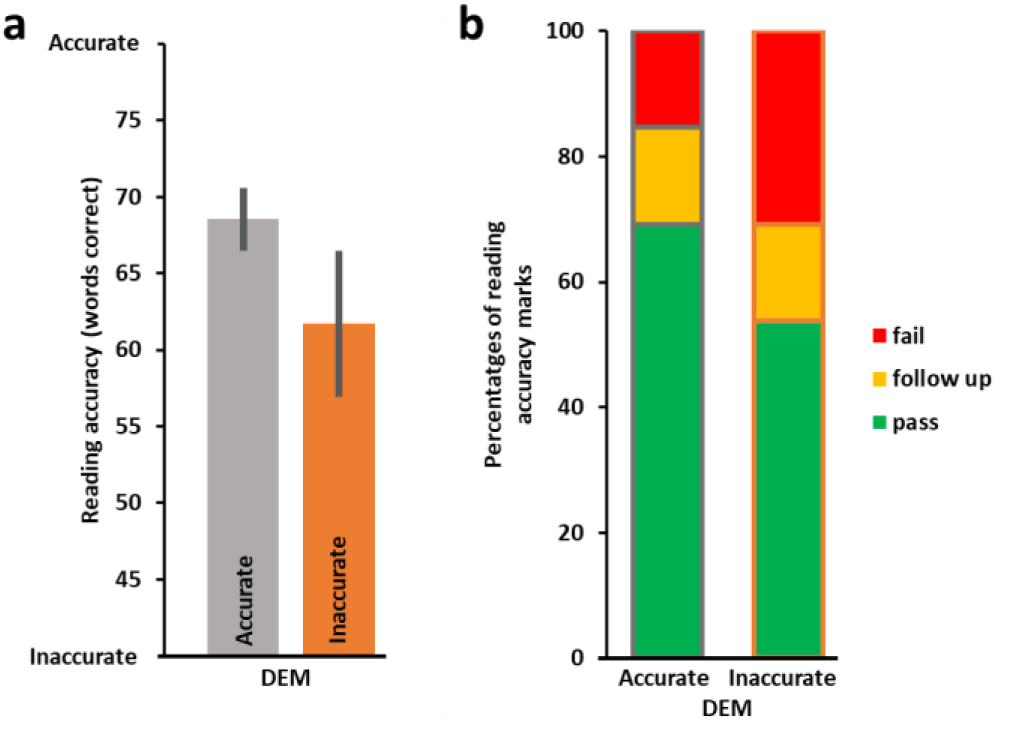
Reading accuracy according to DEM accuracy (n=26). **(A)** Average number of words read correctly (out of 77) by the more accurate (grey, n=13, left) vs the less accurate (orange, n=13, right) DEM performers was not significantly different (*t*(16.264)=-1.31, *p*=.104, 1-tailed). **(B)** Proportions of reading accuracy categorization scores (pass in green, follow up in yellow, fail in red) by DEM accuracy performance. These data tend to suggest that the more accurate DEM group (left column) may receive better reading grades.

To investigate the relationship between the overall DEM categorization (related to the ratio, see Methods) and reading performance, we first examined the distribution of the DEM classification types amongst all the children in the study (see Figure 3) and found that ∼29% were categorized by the DEM as having OMD (DEM Types 2 and 4) while a third were categorized as having normal oculomotor function (Type 1). We then tested whether children categorized as having OMD (Types 2 and 4, n = 8) had worse reading performance than the children categorized by the DEM as having normal oculomotor function (Type 1, n = 9). Reading speed was slower in the children categorized as having OMD (DEM Type 2 and 4 groups, n=8) relative to the children categorized as having normal oculomotor function (DEM Type 1 group, n=9) but this difference was only marginally significant (*t*(9.54) = -1.5, *p*=.083, 1-tailed, see Fig. 4A). Reading accuracy was also worse in the children categorized as having OMD (DEM Type 2 and 4 groups, n=8) relative to the children categorized as having normal oculomotor function (DEM Type 1 group, n=9) but this difference was only marginally significant (*t*(7.61) =1.78, *p*=.057, 1-tailed; see Fig. 4C). In the normal oculomotor group for both reading speed and reading accuracy the proportions of ‘pass’ grades were high (∼80%) and there were no ‘fail’ grades (presented in red in Fig. 4B and D). In contrast, for the OMD groups between a quarter to 50% failed (see Fig. 4B and 4D). Six participants were categorized as Type 3 (language automaticity difficulties) and four participants did not fit into any of the four DEM categorization types and were categorized as an unknown type.

**Fig. 3.**
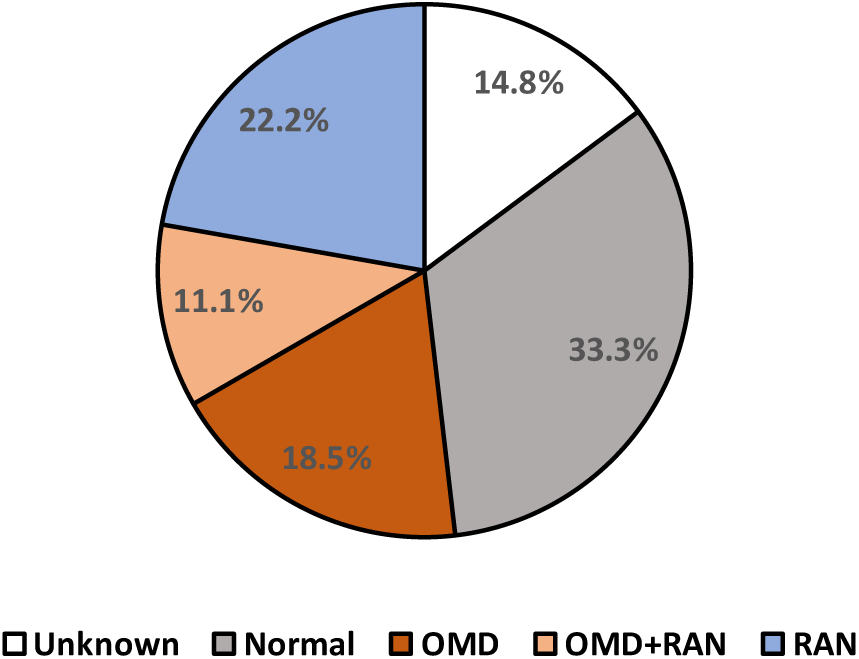
Percentages of children found in each DEM classification type (n=27). The grey section (33.3%, n=9) represents normal language automaticity and oculomotor function (DEM Type 1). Orange shades represent DEM types associated with OMD (29.6%, n=8) with darker orange (18.5%, n=5) representing children categorized by DEM as having normal language automaticity with oculomotor dysfunction (Type 2) and lighter orange section (11.1%, n=3) representing DEM performance categorized as a combination of both oculomotor and language automaticity dysfunctions (Type 4). The blue section (22.2%, n=6) represents abnormal language automaticity function but normal oculomotor function (Type 3). The white section (14.8%, n=4) represents DEM performance that cannot be categorized according to the DEM protocols, possibly due to other visual factors such as accommodative or binocular dysfunctions.

**Fig. 4.**
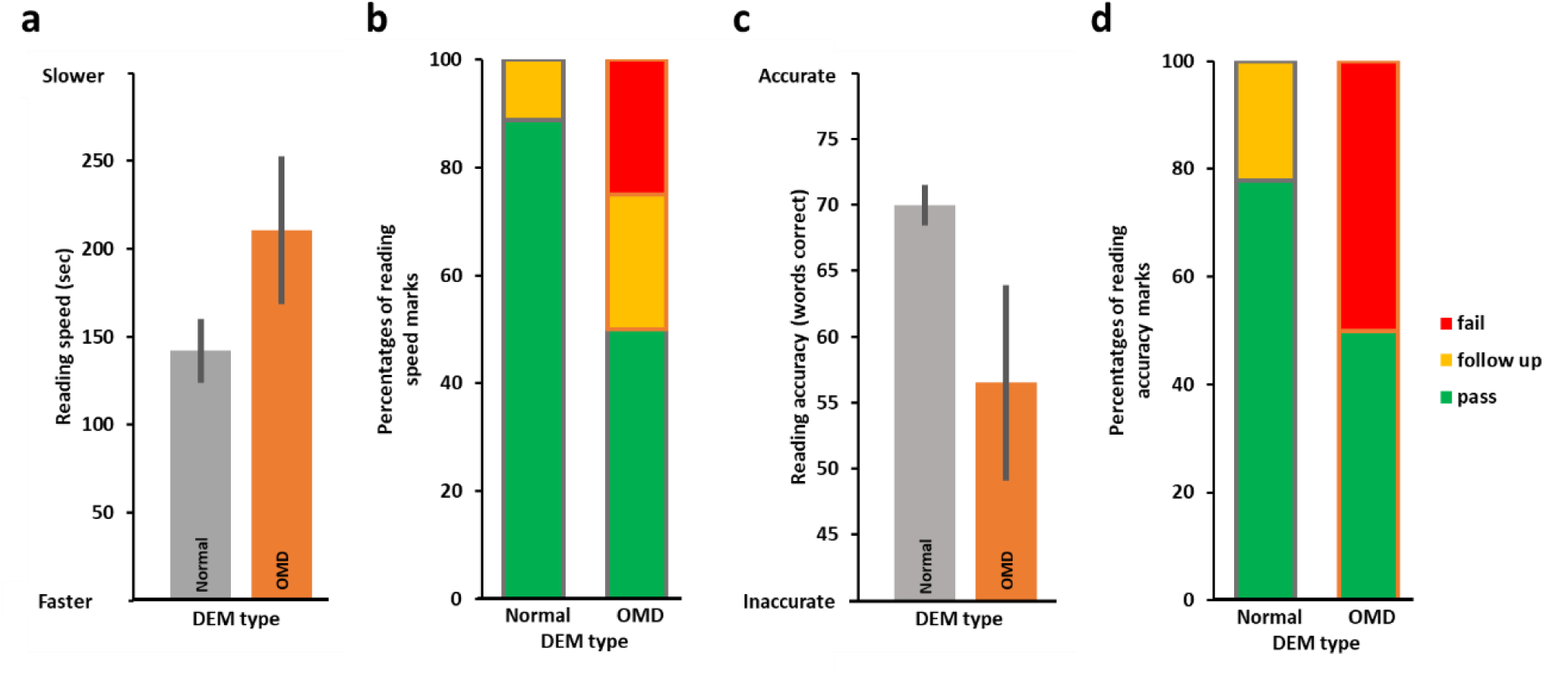
Reading speed and accuracy performance according to DEM Type categorization (n=17). **(A)** Average reading speed in seconds as performed by the normal DEM group (grey Type 1 on left, n=9) and the OMD DEM groups (orange, Types 2&4 on right, n=8). As can be seen the normal DEM group was on average faster in their reading performance (marginally significant, see Results). **(B)** Proportions of reading speed categorization (pass in green, follow up in yellow or fail in red) for the normal DEM group on left and the OMD DEM groups on the right. **(C, D)** Same analysis as in (A, B) but for reading accuracy. Note that there was no fail for either reading speed or accuracy in the normal (Type 1) DEM group.

In summary, the DEM speed results for both vertical and horizontal tests were significantly predictive of reading speed. DEM type was marginally predictive of both reading accuracy and of reading speed. DEM higher accuracy was also on average associated with higher reading accuracy but this was not statistically significant.

### 5.2 Oculomotor evaluation by NSUCO and the relation to reading performance

In addition to the assessment of oculomotor skills by the DEM, oculomotor skills were also assessed independently by the well-established optometric NSUCO oculomotor test. The NSUCO test assesses accuracy of both pursuits and saccades, and we hypothesized that since reading predominantly involves static stimuli and saccades, these would be related to reading accuracy. NSCUO scores four aspects of saccadic performance: ability (typically at ceiling performance), accuracy (ability to refrain from overshooting or undershooting when trying to reach targets), head movements and body movements. As body movements accompanying saccades usually involve head movements as well (but the opposite is not always the case) we examined saccadic head movement scores and their relationship to reading ability. When we split the class according to pass/fail criterion for the saccadic head movement sub-score, we found the NSUCO performance was not significantly predictive of reading accuracy (*t*(9.039)=-1.05, *p*=.160, 1-tailed). As can be seen in Fig. 5A, the group that did not succeed on the NSUCO saccadic head movement performance (n=12) performed on average less accurately on the reading accuracy test than the group that achieved age-appropriate scores on the NSUCO saccadic head movement performance (n=15). We then compared proportions of ‘pass’, ‘requires follow up’ and ‘fail’ grades in the reading accuracy test for the NSUCO age-appropriate head movement group (n=15) and for the below age-appropriate group (n=12). The proportion of reading accuracy ‘pass’ grades in the NSUCO age-appropriate head movement group was 61.1% and 16.7% with a ‘fail’. The below age-appropriate NSUCO group had a 56% ‘pass’ rate for reading accuracy and 33.3% ‘fail’ (Fig. 5B).

**Fig. 5.**
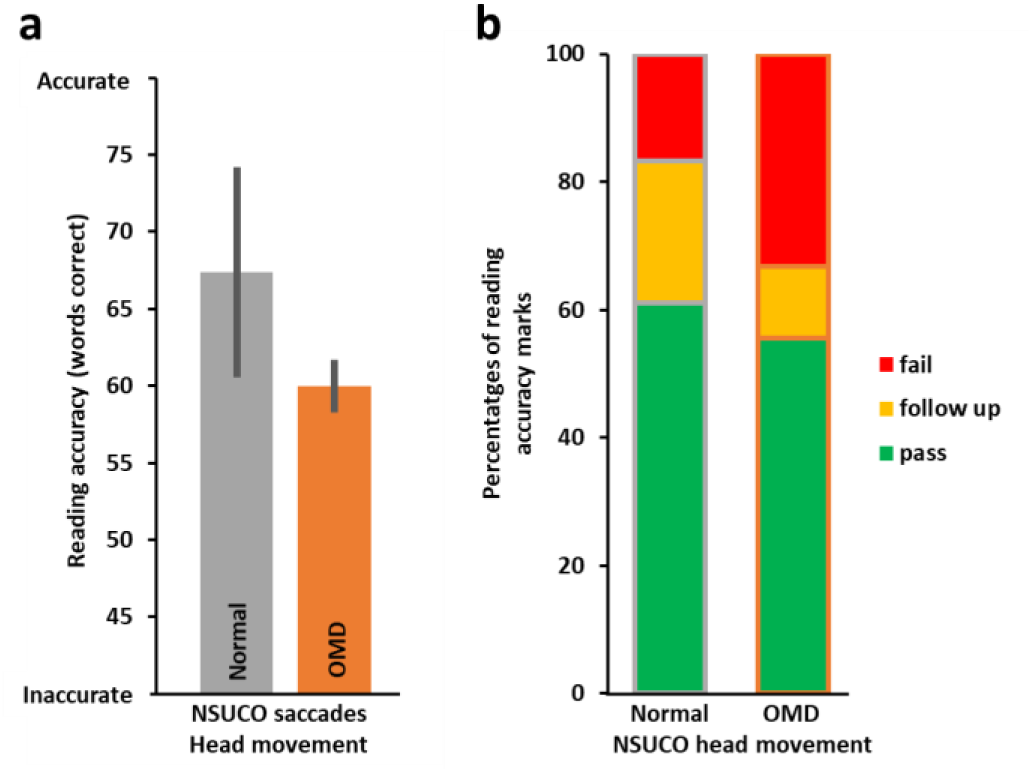
Reading accuracy performance according to standardized optometric NSUCO subscore that grades the amount of head movement resulting from saccades (n=27). **(A)** Average number of words read correctly as performed by the age-appropriate NSUCO head movement group (grey, n=15) and below age-appropriate (orange) group (n=12). The age-appropriate group was more accurate on average in their reading. (B) Proportions of reading accuracy categorization (‘pass’, ‘follow up’ or ‘fail’) for both groups.

### 5.3 Distance visual acuity (VA) and the relation to reading performance

As control, we also assessed whether non oculomotor visual measures may be associated with reading acquisition performance. We compared distance VA, a commonly used measure to assess vision in school-aged children, with reading performance (speed and accuracy). Surprisingly when we correlated dVA with reading speed, we found that poorer dVA was marginally predictive of faster reading (*r_s_*(24)=.37, *p*=.062) as well as being significantly predictive of more accurate reading *(r_s_*(24)=-.47*, p*=.014). When comparing the reading of the better dVA group with the poorer dVA group we found that children with worse dVA (less than 6/6, n=10) were significantly faster in their reading performance (*t*(23.284)=-2.22, *p*=.037, 2-tailed; Fig. 6A) and significantly more accurate in their reading (*t*(17.363)=2.82, *p*=.012, 2-tailed) than children with normal dVA (6/6, n=16, see Fig. 6B). Amongst children with normal dVA (i.e., 6/6) the proportions of ‘pass’ for the reading speed and accuracy were ***lower*** than in the group of reduced dVA (i.e., less than 6/6). There were no ‘fail’ in the reduced dVA group while in the normal dVA group there were >30% ‘fail’ marks (Fig. 6C, D). It is important to note that many of the participants with reduced dVA (less than 6/6) had only marginally reduced dVA. Six children with poorer dVA also had uncorrected myopia and/or astigmatism. Our results indicate that dVA, which we used as a control, was not a good predictor of reading performance from close distance.

**Fig. 6.**
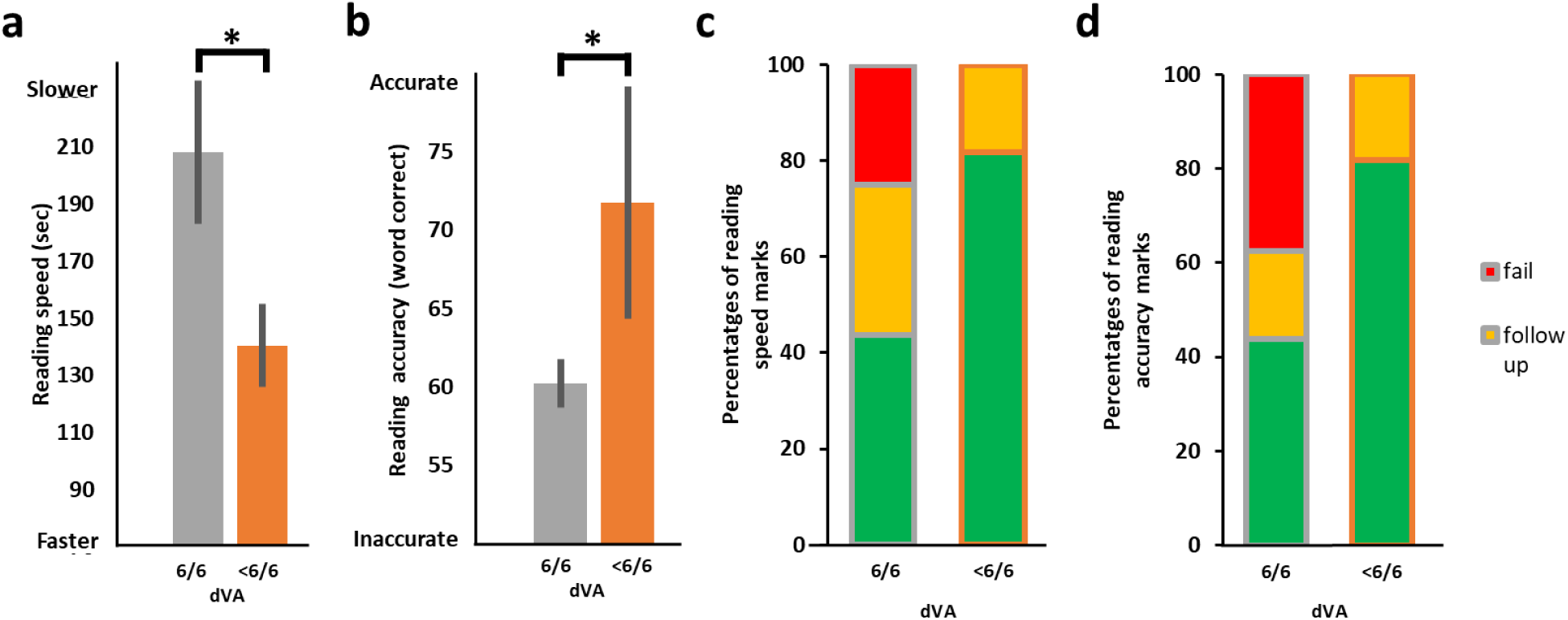
Reading performance in children with better and poorer VA (n=26). **(A)** The Y-axis represents reading time in seconds. The right orange bar depicts reading speed (in words per minutes) of the below-average dVA (<6/6) group (n=10), the left grey column that of the average dVA (6/6) group (n=16). Reading speed of the better acuity group was significantly *slower* than that of the poorer acuity group (*t*(23.284)=-2.22, *p*=.037, 2-tailed). **(B)** Similar to (A) with Y-axis representing the number of words read accurately out of the 77 words in the test. Reading accuracy performance of the better acuity group was marginally less accurate than that of the group with less than average (<6/6) dVA (*t*(17.363)=2.82, *p*=.012, 2- tailed). **(C)** The stacked bar chart displays proportions of reading speed categorization (pass in green, follow up in yellow, fail in red), right bar - reading score distribution of the less than average dVA group, left bar - similarly for the normal dVA. Note that there were no ‘fails’ in the <6/6 group and the ‘pass’ rate was higher than in the 6/6 group. **(D)** Same analysis as in (C) but for reading accuracy.

### 5.4 Linear regression analyses

To further corroborate our findings, we also ran linear regression analyses assessing whether aspects of the DEM and/or dVA were predictive of reading accuracy or speed. As can be seen in Table 1 (top part, reading accuracy), dVA (Model 1) on its own was not significantly predictive of reading accuracy but DEM horizontal accuracy was (see Model 2) and so was the DEM ratio (Model 3). We also tested whether all predictors combined (dVA, DEM accuracy and DEM ratio) may better explain reading accuracy (Model 4) but this model did not outperform Model 3 based on the DEM ratio alone (compare Model 3 with *R*^2^ of 0.2394 vs Model 4 with *R*^2^ of 0.2122). Reading speed analyses (Table 1 lower part) revealed that dVA alone (Model 1) was also not significantly predictive of reading speed but DEM horizontal speed (Model 2) or DEM vertical speed (Model 3) on their own were significant. Importantly, the model that outperformed all models in explaining reading speed was Model 4 that combined dVA, DEM horizontal and vertical speed which were all significant with *R*^2^ of 0.4353. This indicates that the combination of both dVA and DEM speed measures gave the best prediction for reading speed.

**Table 1.** Linear regression analyses results. Top – models explaining reading accuracy, bottom – models explaining reading speed. Each row specifies one predictor (bottom row the overall model estimation summary), each column details the outcome of one linear model with model results included only in cells that were part of that model. In each cell, the model estimate for that variable ± the standard error and in parenthesis its significance. Significant values (apart from the intercept) indicated in bold, asterisks indicate predictor significance (* for *p*<.05, ** for *p*<0.01).

| READING ACCURACY |  |  |  |  |
| --- | --- | --- | --- | --- |
|  | Model 1 | Model 2 | Model 3 | Model 4 |
| INTERCEPT | 75.43 ± 9.05<br>(1.52*10<sup>-8***</sup>) | 21.9304 ± 17.3171<br>(.2175) | 58.74729 ± 3.02448<br>(3.47*10<sup>-16***</sup>) | 48.62904 ± 22.55831<br>(.0423*) |
| DVA | -12.60 ± 10.11<br>(.224) | - | - | -7.13391 ± 9.32139<br>(.4522) |
| DEM HORIZONTAL ACCURACY | - | **0.4609 ± 0.1852**<br>**(.0202\*)** | - | 0.19512 ± 0.24644<br>(0.4370) |
| DEM RATIO | - | - | **0.08743 ± 0.02936**<br>**(.00654\*\*)** | 0.06011 ± 0.04151<br>(0.1618) |
| OVERALL MODEL ESTIMATION | *F*(1,24)=1.556,<br>*p*=.2243,<br>Adjusted *R*<sup>2</sup> = 0.02174 | *F*(1,24)=6.194<br>*p*=.02015<br>Adjusted *R*<sup>2</sup> = 0.172 | *F*(1,24)=8.867<br>*p*=.006543<br>Adjusted *R*<sup>2</sup> = 0.2394 | *F*(3,22)=3.245<br>*p*=.0414<br>Adjusted *R*<sup>2</sup> = 0.2122 |
| READING SPEED |  |  |  |  |
|  | Model 1 | Model 2 | Model 3 | Model 4 |
| INTERCEPT | 91.0 ± 59.21<br>(.137) | 212.2101 ± 17.9623<br>(1.73*10<sup>-11***</sup>) | 304.3051 ± 57.7368<br>(2.1*10<sup>-5 ***</sup>) | 232.7528 ± 58.8213<br>(.00067***) |
| DVA | 107.05 ± 66.12<br>(.119) | - | - | **117.2505 ± 53.8036**<br>**(.04030\*)** |
| DEM HORIZONTAL SPEED | - | **-0.5290 ± 0.1724**<br>**(.00526\*\*)** | - | **-1.5051 ± 0.5509**<br>**(.01216\*)** |
| DEM VERTICAL SPEED | - | - | **-1.4333 ± 0.6532**<br>**(.0381\*)** | **-0.4125 ± 0.1545**<br>**(.01400\*)** |
| OVERALL MODEL ESTIMATION | *F*(1,24)=2.621,<br>*p*=.1185,<br>Adjusted *R*<sup>2</sup> = 0.06089 | *F*(1,24)=9.419,<br>*p*=.005264,<br>Adjusted *R*<sup>2</sup> = 0.2519 | *F*(1,24)=4.815,<br>*p*=.03813,<br>Adjusted *R*<sup>2</sup> = 0.1324 | *F*(3,22)=7.424,<br>*p*=.001301,<br>Adjusted *R*<sup>2</sup> = 0.4353 |

## 6 Discussion

In this study, we retrospectively investigated the relationship between oculomotor performance and reading acquisition in a normative class of Hebrew speaking first graders. We found that different scores of a well-established test for oculomotor skills, the DEM, were significantly predictive of Hebrew reading performance. For an additional test routinely used in the optometry clinic, the NSUCO, the head movement sub-score of the test was on average indicative of early Hebrew reading acquisition ability but this was not significant. Interestingly, better distance visual acuity was not positively associated with better reading performance.

Many studies have assessed reading abilities in different developmental conditions (ASD (33), attention related disorders (34), developmental dyslexia (22,35), APD (36)). Here we investigated the relation between reading acquisition and developmental visual skills focusing on the general population (normative class within the state educational system) rather than on specific developmental subgroups. We found a clear relationship between oculomotor-related measures and reading acquisition, and the prevalence of OMD (as estimated by the DEM) was ∼29% which is in line with earlier estimates of 15-24% (5,6) prevalence in good readers and much higher prevalence in poor readers (5,7). While our investigation was of a normative class in the general population, we cannot rule out the possibility that some of the children may have had reading-related or attention-related difficulties alone or in addition to visual issues, given the prevalence of these conditions in the general population (5-10% (37) and roughly 5% (38), respectively). It is unclear if the proportion of OMD in the class we examined truly reflects OMD prevalence in the entire population and a bigger cohort would be needed to further substantiate these overall estimates. However, even if the prevalence of OMD is found to be smaller, even halved, this would suggest that a substantial portion of children entering the educational system may have oculomotor-related difficulties which are likely to be related to reading acquisition difficulties, as our results indicate. It would therefore be beneficial for children who exhibit reading difficulties to consider including testing for OMD in addition to the current tests.

Previous studies that have investigated the relationship between oculomotor skills and reading abilities examined this at later stages of reading acquisition (22,35,39,40) and often the focus was on groups of children with reading related difficulties (e.g. Refs. (22,35)). For example, a review by Kulp and Schmidt suggests a correlation between oculomotor dysfunction and reading disability across different age groups in an English reading population, and these are based on different oculomotor control assessments than those employed in our study (41). Raghuram et al. found in a group of developmental dyslexic English readers aged 8-11 years (middle-end of elementary school) that their oculomotor skills as assessed by the DEM were significantly lower than those of typically developing controls (22). In our study we examined the relationship between oculomotor function and reading in a normative class at the initial stages of reading acquisition (1^st^ grade, 6-7 years of age) in Hebrew readers (read from right to left) when two different substantiated oculomotor tests (DEM (26,30–32,42) and NSUCO (42–47)) were employed. Despite differences between our study and earlier ones we still found that oculomotor skills were associated with reading acquisition. Due to the small cohort size, future replications will be necessary, however these results suggest that oculomotor function may be related to reading across languages (regardless of orthographic depth or reading direction). In contrast to previous studies in which reading and NSUCO scores were shown to be significantly correlated, our NSUCO findings were weaker than the DEM measures were (see Results and Figures 1, 2 and 4) and this may be due to our small cohort size relative to earlier studies with larger cohorts (45,47). Furthermore, previous studies investigating the performance of NSUCO and reading ability were performed with English reading children whereas our study used Hebrew reading that uses a different (right-to-left) alphabet. Additionally, during the initial reading acquisition of Hebrew, diacritical markers are used to help novice readers pronounce the words correctly which is very different from English reading acquisition that does not use diacritical markers and instead depends on contextual cues for reading accuracy and correct pronunciation. We hypothesize that the visual demands for initial stages of Hebrew reading may therefore have different oculomotor demands than those demanded during English reading acquisition. Therefore, the NSUCO test which has been shown to be an effective nonverbal assessment tool for evaluating gross ocular motor function, may not be sensitive enough to predict reading ability in young Hebrew readers.

In the literature, the relationship between dVA and reading is still unclear. In many countries including European countries (14) and in the USA, visual evaluation in schools often includes only distance visual acuity (dVA) which is effective in detecting amblyopia. A leading cause of reduced dVA is short-sightedness (myopia), and its prevalence in the age range of our cohort is lower than that of far-sightedness (hyperopia)(48,49). Bruce et al. (50) report that dVA is associated with developing literacy skills at the age of 4-5. In their study they used a subtest of the Woodcock Reading Mastery Tests-Revised (WRMTS-R) that demands letter recognition and not whole word or sentence reading, perhaps due to the young children’s pre-reading age^51^. Such visually local letter recognition may rely on similar functions as VA testing, which may be one of the factors contributing to the correlation they found. A recent review by Hopkins et al. on dVA and reading ability concludes that the impact of dVA on reading ability is still unclear (51) and there are other studies reporting that dVA is not associated with reading ability. For example, one longitudinal study assessing 1143 nine- and ten-year-olds found that there was no correlation between dVA and academic school performance (52). However, they did not directly assess the relationship between dVA and reading. Additionally, as the age of the children in their study was 9 and 10, it is difficult to draw any inferences from these findings about reading acquisition that typically occurs at earlier age. Another study also found that dVA was not predictive of reading skills (53). In our study we also found that dVA was not positively associated with reading acquisition, and our results may be explained by the fact that the reading examination was performed at close viewing distance using large font sizes while dVA was performed from a far viewing distance. While dVA is important for multiple school-related purposes (seeing the board, reading the teacher’s lips, social communication), the jury is still out about the contribution of distance VA to reading (see also Ref. (54)). Nevertheless, in light of our results, we suggest that children who are found to be struggling with reading should be referred to a comprehensive vision evaluation that includes oculomotor assessments ^55,56^. Detection of OMD-related difficulties in early readers could allow consultation with specialists to consider dedicated visual support for improving scholastic performance (12,55–60) across school-aged children.

## 7 Conclusion

OMD prevalence in school-aged good readers has been suggested to be approximately 15-24% (5,6,22) compared to much higher estimates in children diagnosed with a reading disability (5,7). Here, retrospectively assessing optometric evaluation results of a normative first grade class, we found that ∼29% of the children were categorized by the DEM as having OMD-related difficulties. While future assessments of the relationship between OMD-related measures and reading performance are necessary with larger cohort sizes, our study suggests that reading difficulties from near distances (approx. up to 40cm) among Hebrew early readers are significantly associated with oculomotor difficulties (i.e. OMD). We propose that adding oculomotor evaluations for school-aged children who are struggling with reading may be beneficial for enabling accurate and fluent reading.

## 11 Author Contributions

SGD: Conceptualization, Data curation, Formal analysis, Funding acquisition, Investigation, Methodology, Project administration, Resources, Supervision, Validation, Visualization, Writing – original draft, Writing – review & editing. AP: Conceptualization, Data curation, Formal analysis, Investigation, Methodology, Project administration, Resources, Validation, Visualization, Writing – original draft, Writing – review & editing.

## 13 Institutional Review Board Statement

The study was conducted in accordance with the Declaration of Helsinki and approved by the Bar-Ilan University Interdisciplinary Studies Unit Research Ethics Committee (date of approval, 1 January 2019).

## 14 Informed Consent Statement

Participation consent was waived as this study was retrospective based on data stripped of identifying information.

## 15 Acknowledgments

We thank Olga Kreichman for her comments on the manuscript, and Racheli Rubinov for her assistance in collecting and assessing the reading ability of the children.

## 16 Data Availability Statement

The dataset related to this study can be found in the OSF repository at https://osf.io/shw2v/.

